# Polysialylated NCAM distinguishes differentiated myoblasts from proliferating myoblasts prior to fusion

**DOI:** 10.1101/2025.04.08.647904

**Authors:** Tetsutaro Kikuchi, Tatsuya Shimizu

## Abstract

The differentiation of skeletal muscle myoblasts is a crucial step in muscle regeneration and hypertrophy. However, the phenotypic changes that occur in myoblasts just before myotube formation have remained poorly understood. In this study, we established a novel flow cytometric method to examine these changes. We identified two distinct populations in human desmin-positive myoblasts based on immunological NCAM (neural cell adhesion molecule) staining intensity after paraformaldehyde fixation. Our results indicate that these populations correspond to differentiated and proliferating myoblast phenotypes. Further analyses revealed that antigenicity for a polysialylated-NCAM (PSA-NCAM) antibody is strongly correlated with this fixation-resistant antigenicity of NCAM, suggesting that polysialylation is associated with myogenic differentiation. Using magnetic-activated cell sorting (MACS), we separated PSA-NCAM positive and negative myoblast fractions and demonstrated that PSA-NCAM positive cells show higher differentiation marker expression and faster myotube formation. These findings suggest that PSA-NCAM can serve as a valuable surface marker to distinguish between proliferating and differentiated myoblasts. Our method provides new insights into the phenotypic heterogeneity of myoblasts and offers an approach to studying the early stages of muscle cell differentiation before myotube formation.

## Introduction

Skeletal muscle responds to mechanical loading or injury by activating muscle satellite cells, which regenerate or strengthen the tissue (Bischoff, 1986; Konigsberg et al., 1975; MAURO, 1961). In this process, intermediate myogenic precursor cells are commonly referred to as ‘myoblasts’, which includes the two phenotypes: ‘proliferating myoblasts’ and ‘differentiated myoblasts’, where the latter is sometimes called ‘myocytes’ (Hindi et al., 2013; Iberite et al., 2022; Mierzejewski et al., 2020; Pajalunga and Crescenzi, 2021). With advances in single-cell RNA sequencing technology, detailed analysis of cellular populations within human skeletal muscle samples has become increasingly feasible (De Micheli et al., 2020; Fitzgerald et al., 2023). However, the cell populations involved in muscle regeneration and hypertrophy are rare against the overall muscle tissue, necessitating continued use of in vitro experiments for detailed mechanistic analyses.

In conventional in vitro differentiation induction experiments, low-serum media have been used to promote myotube formation. The drawback of this approach is causing cell fusion to occur simultaneously with differentiation, making it difficult to precisely evaluate phenotypic changes prior to fusion (Katayama et al., 2023; Wang et al., 2023). Therefore, establishing methodology to investigate this myoblasts’ phenotypic changes in the pre-fusion period should be valuable.

In this regard, we observed that cultured human skeletal muscle myoblasts (that are positive for desmin) could be divided into two populations based on their staining intensities with the NCAM (neural cell adhesion molecule) antibody. This led us to hypothesize that these two groups correspond to the proliferating and differentiated phenotypes of myoblasts.

Since myoblasts initiate myotube formation within 24 hours of switching to differentiation media, we plated cells at extremely low density to prevent cell fusion, thereby allowing flow cytometric analysis. The results supported our initial hypothesis. Furthermore, we found that an antibody against polysialylated NCAM functions as a potential surface marker distinguishing between the proliferating and differentiated myoblast phenotypes.

Methods to investigate the differentiation state of myoblasts without inducing myotube formation have not been established previously, and we believe that our approach represents a valuable tool for studying the differentiation process of skeletal muscle myoblasts in detail, especially just before the fusion.

## Results

### Fixation resistant antigenic NCAM (fra-NCAM) divides desmin-positive cells into two groups

In investigating immunostaining conditions for NCAM and desmin (markers for myoblasts) in human skeletal muscle cells, we observed that the staining intensity of NCAM antibody (clone: MEM-188) significantly decreased after fixation with paraformaldehyde (Fig. 1a). While it is well known that fixation can reduce antigenicity for some antibodies, we surprisingly found that, within the desmin-positive cell population, two distinct groups were evident based on NCAM staining intensities after fixation in certain muscle-derived cell populations (Fig. 1b). This observation suggests the presence of phenotypic heterogeneity regarding the expression state of NCAM in desmin-positive cells. We termed this fixation-resistant antigenic NCAM as ‘fra-NCAM’

**Fig. 1.**
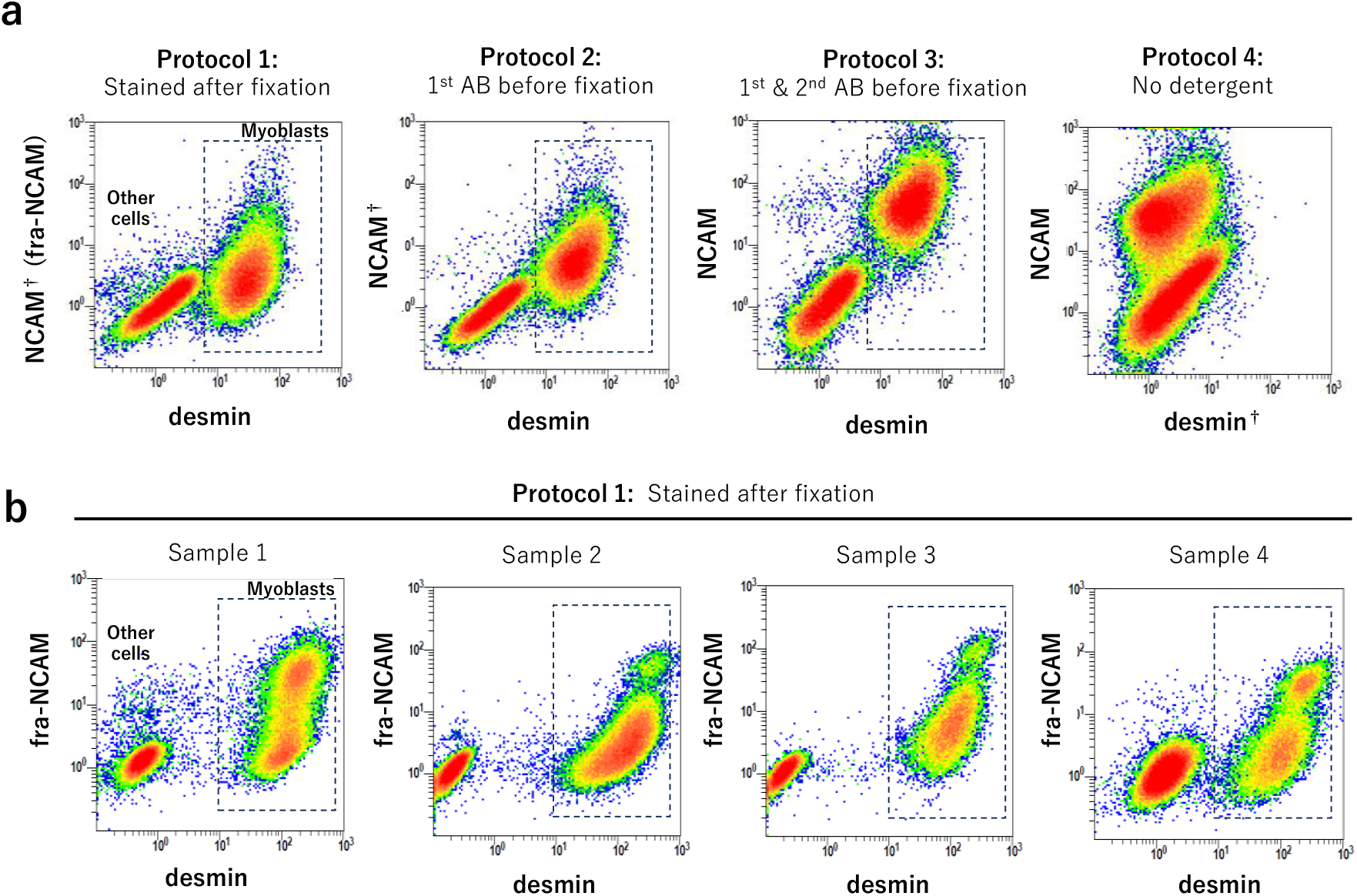
Fixation resistant antigenic NCAM (fra-NCAM) divides desmin-positive cells into two groups. a) Flow cytometric analysis for human skeletal muscle cells stained for NCAM and desmin by four different protocols. b) Four samples of human skeletal muscle cells were stained by Protocol 1. Daggers (†) mark non-standard staining condition. fra-NCAM indicates the intensity for NCAM in Protocol 1.

### Effect of growth factors on fra-NCAM intensity

Given that desmin-positive cells are thought to be myogenic, it is speculated that these two phenotypes may correspond to proliferating and differentiated myoblasts. To explore this idea, we investigate how fra-NCAM intensity is associated with myoblast differentiation. First, we conducted differentiation induction culture for human skeletal muscle cells (hSkMCs). hSkMCs were cultured in serum-free medium (Neurobasal + B27 + GlutaMAX) for three days at a low density that reduces cell fusion and analyzed by flow cytometry. The ratio of fra-NCAM bright cells increased after the induction (Fig. 2a).

**Fig. 2.**
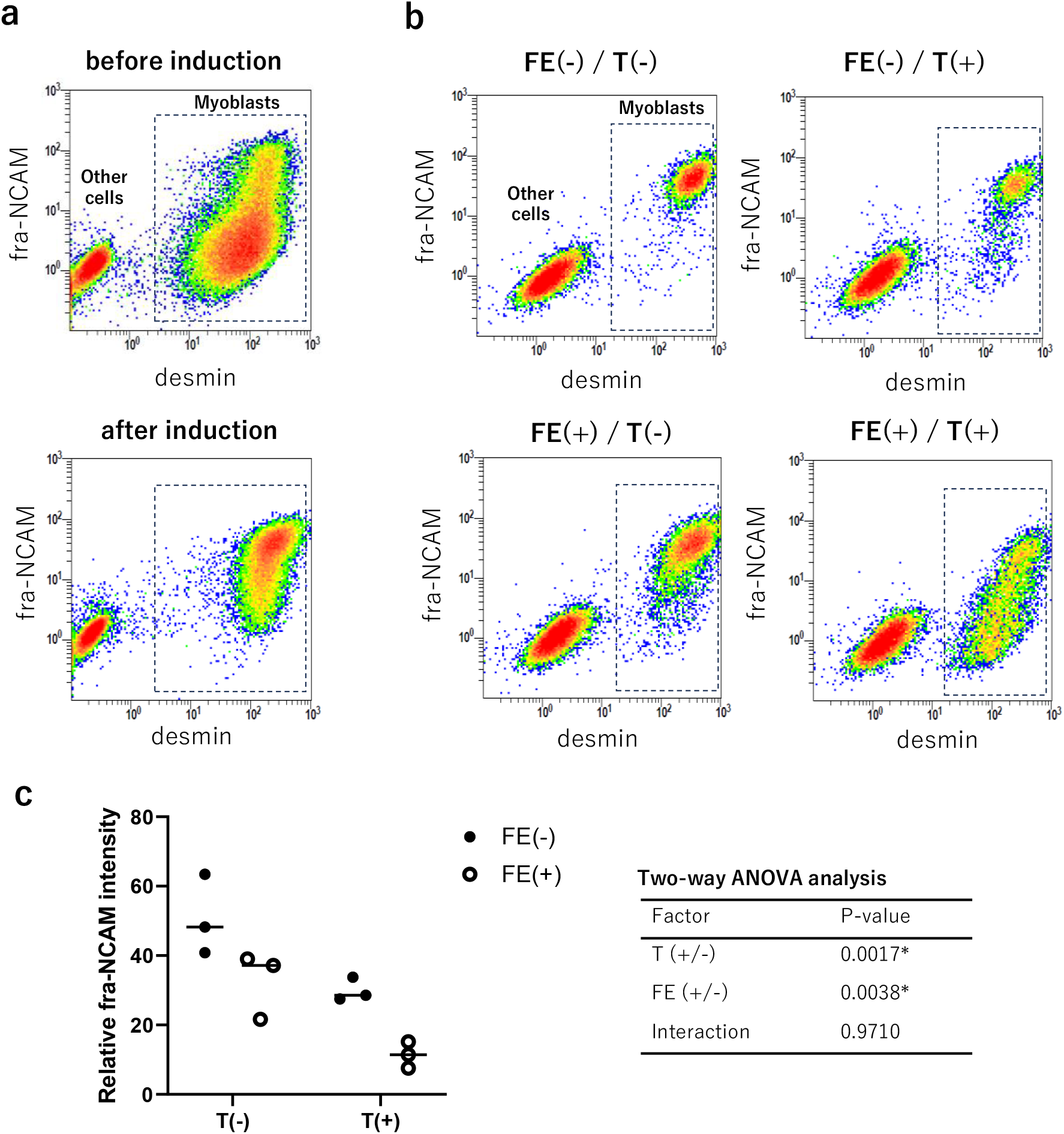
Effect of growth factors on fra-NCAM. a) hSkMCs were cultured in serum-free medium (Neurobasal+ B27+GlutaMAX) for three days. b) hSkMCs were cultured in serum-free medium (DMEM+ITS+rHSA+iMatrix-221) with or without growth factors. c) Statistical analysis for the effect of growth factors against fra-NCAM intensity in desmin-positive cells. FE: FGF-2 (10 ng/mL) + EGF (10 ng/mL). T: TGF-β1 (1 ng/mL). *: p < 0.05.

Next, we examined the effect of growth factors known to attenuate myoblast differentiation (Fig. 2b) (Brunetti and Goldfine, 1990; Florini et al., 1986; Leroy et al., 2013; Weyman and Wolfman, 1998; Wicik et al., 2010). To minimize the influence of serum and other growth factors, we used a simpler serum-free medium (DMEM + ITS + recombinant HSA + iMatrix-221). hSkMCs cultured in serum-free medium for three days displayed high fra-NCAM intensity. The combination of FGF-2+EGF, as well as TGF-β1, reduced the intensity of fra-NCAM in desmin-positive cells. The addition of both FGF-2+EGF and TGF-β1 resulted in an even greater reduction in fra-NCAM intensity (Fig. 2c). These results suggest that the fra-NCAM bright group corresponds to differentiated myoblasts, while the fra-NCAM dim group corresponds to proliferating myoblasts.

### Effect of signal inhibitors on fra-NCAM

To further verify the characteristics of the fra-NCAM bright and dim groups, we examined the effects of known signaling inhibitors (Fig. 3). The addition of SB505124, an inhibitor of TGF-β signaling, showed a significant increase in fra-NCAM intensity under growth factor-supplemented conditions. Similarly, U0126, an ERK1/2 signaling inhibitor, led to a significant increase in fra-NCAM intensity in the presence of growth factors, while PD184352, another ERK1/2 inhibitor, did not show significant effects. LY-294002, an inhibitor of PI3K/Akt signaling, reduced fra-NCAM intensity irrespective of the presence of growth factors. For inhibitors targeting p38 signaling, SB203580 and BIRB796, as well as SP600125 for JNK signaling, only BIRB796 significantly reduced fra-NCAM intensity in the presence of growth factors, while no significant differences were observed for the other conditions. Representative scatter plots are shown in Fig. S1.

**Fig. 3.**
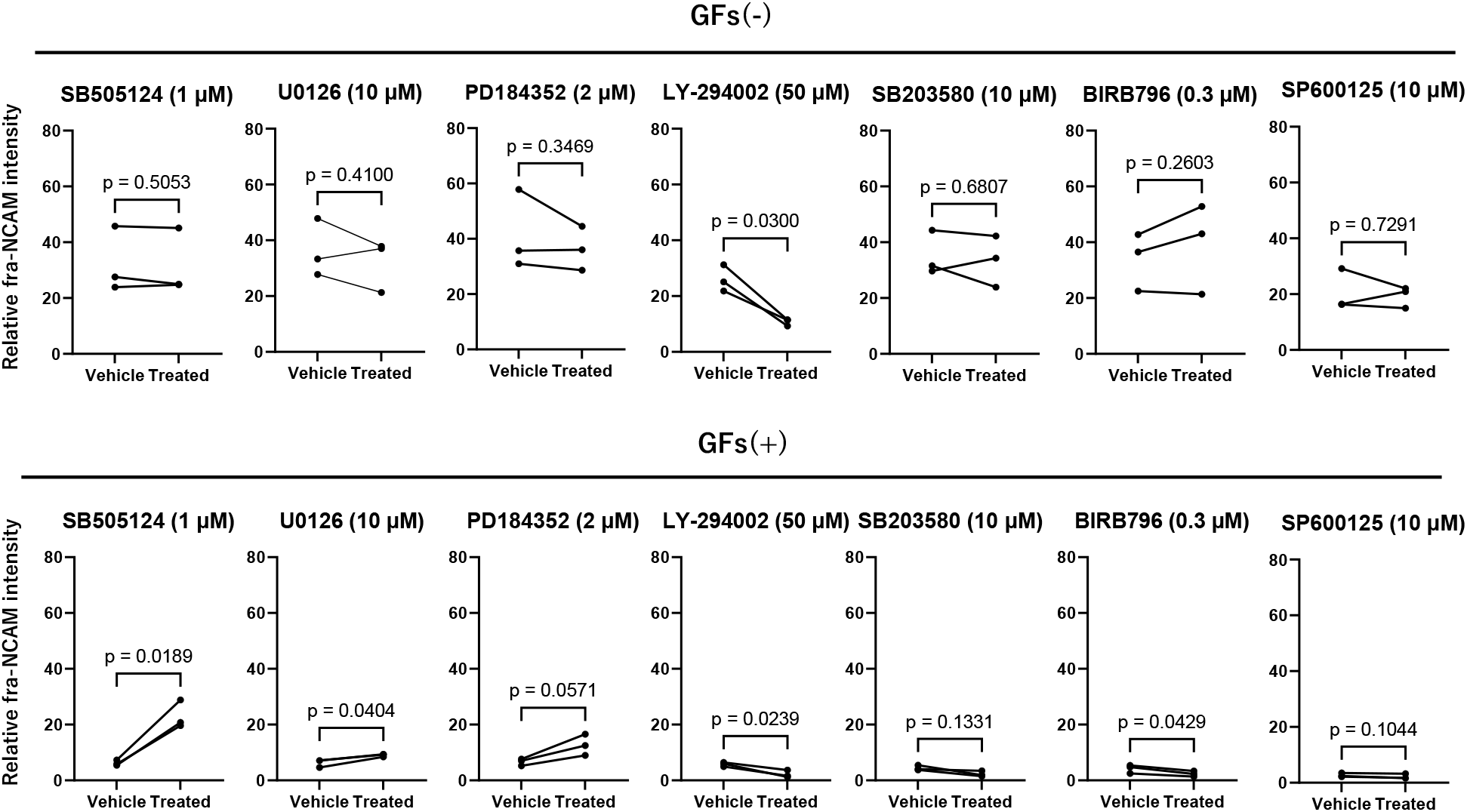
Effect of signal inhibitors on fra-NCAM. hSkMCs from three different strains were cultured in serum-free medium for 3 days with or without signal inhibitors under the conditions with or without growth factors (GFs: 10 ng/mL FGF-2, 10 ng/mL EGF and 1 ng/mL TGF-β1).

### Polysialylation may be linked to fixation resistant antigenicity on NCAM

Changes in NCAM antigenicity might reflect molecular modifications or alterations in multimeric complexes. We, therefore, investigated polysialic acid modification of the NCAM molecule, a feature that is well studied in neuronal cells (Brusés et al., 1995; Hildebrandt et al., 2007; Röckle et al., 2008). Using a polysialylated NCAM (PSA-NCAM) specific monoclonal antibody (clone: 12E3) for flow cytometric analysis, we found that PSA-NCAM staining strongly correlated with fra-NCAM (Fig. 4). These results suggest that polysialylation may be associated with the fixation-resistant antigenicity observed in NCAM.

**Fig. 4.**
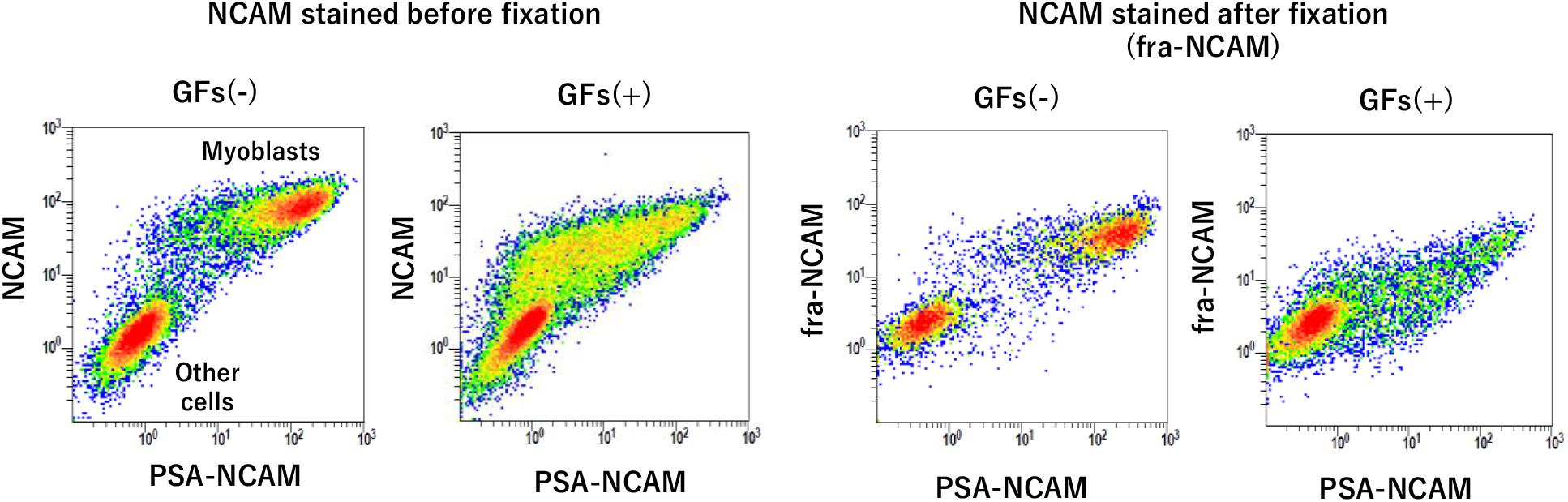
Relation of fixation-resistance of NCAM and polysialylation. hSkMCs were cultured in serum-free medium for 3 days under the conditions with or without growth factors (GFs: 10 ng/mL FGF-2, 10 ng/mL EGF and 1 ng/mL TGF-β1) and stained for NCAM and polysialylated NCAM (PSA-NCAM) antibodies by two different protocols where NCAM antibody was applied before or after fixation.

### PSA-NCAM positive myoblasts presented more differentiated phenotype than PSA-NCAM negative myoblasts

Lastly, we investigated whether PSA-NCAM could serve as a marker to distinguish between proliferating and differentiated myoblasts. Using magnetic-activated cell sorting (MACS) with PSA-NCAM and NCAM antibodies, we were able to separate PSA-NCAM positive and negative myoblast fractions from hSkMCs cultured in commercially available growth medium (SkGM2) (Fig. 5a). Gene expression analysis of these two fractions revealed that MYOG, MYH7, MYH3, TNNT3, TNNI2, and ACTA1 were expressed at higher levels in PSA-NCAM positive cells (Fig. 5b). Moreover, a tube formation assay demonstrated that PSA-NCAM positive cells exhibited a significantly higher proportion of myosin 4 positive cells after both 24 and 48 hours of differentiation compared to PSA-NCAM negative cells (Fig. 5c, 5d).

**Fig. 5.**
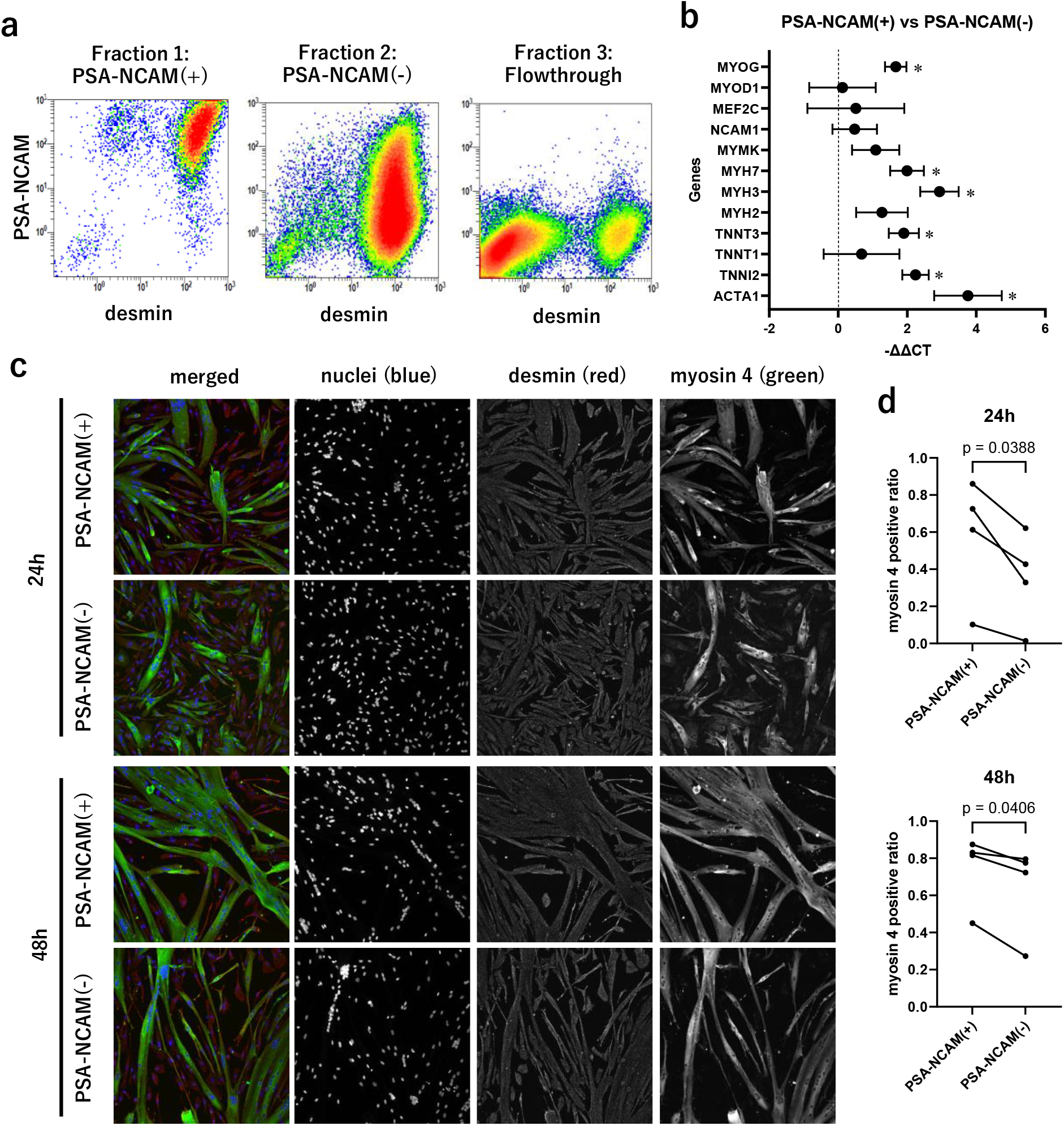
MACS separation exploiting PSA-NCAM and NCAM antibodies. a) hSkMCs were fractioned by two-step MACS using PSA-NCAM and NCAM antibodies. b) Gene expression of PSA-NCAM(+) and PSA-NCAM(-) group of hSkMCs was analyzed by qPCR. Asterisks indicate p < 0.05 in paired Student’s t-test for three cell strains. c) Representative immunofluorescence images in tube-formation assay. Red: F-actin. Green: myosin 4. Blue: nuclei. d) Image-based cytometric analysis for myosin 4 positive ratio. P-values are based on paired Student’s t-test for four cell strains.

## Discussion

In this study, we aimed to analyze phenotypic changes in myoblasts just prior to myotube formation; a stage that has not been thoroughly investigated. To achieve this, we exploited flow cytometric analysis, which has not been widely used because the analysis of fused cells is challenging. Initially, we analyzed the populations of human skeletal muscle myoblasts cultured in commercial growth media and found two distinct groups within the desmin-positive cells that differed in NCAM antibody staining intensity after fixation (Fig. 1). To further investigate how these two groups correspond to skeletal muscle differentiation, we established a low-density culture system using serum-free media to prevent cell fusion and allow quantitative evaluation by flow cytometry.

Our results showed an increase in fra-NCAM bright population after serum-free differentiation induction (Fig. 2a). Additionally, the addition of growth factors, which are reported to attenuate myoblast differentiation, suppressed the increase of the fra-NCAM bright population in the serum-free culture system (Fig. 2b, 2c). Further analysis using signaling inhibitors revealed that the part of inhibitors for TGF-β, ERK1/2, and p38^MAPK^ signaling significantly increased fra-NCAM intensity under growth factor-supplemented conditions, while the PI3K/Akt signaling inhibitor decreased fra-NCAM intensity regardless of the presence of growth factors (Fig. 3). These findings suggest that fra-NCAM becomes positive as myoblasts differentiate.

The mechanism responsible for the changes in NCAM antibody staining after fixation remains unclear; however, we found that the staining intensity of fra-NCAM strongly correlated with the antigenicity for PSA-NCAM antibody (Fig. 4). Polysialylation of the NCAM molecule has been extensively studied in neuronal cells (Brusés et al., 1995; Hildebrandt et al., 2007; Röckle et al., 2008), with some of its physiological significance already elucidated, but there are no precise reports regarding the polysialylation of NCAM in skeletal muscle myoblasts. Although this study does not provide any information on the physiological function of PSA-NCAM, our results suggest that PSA-NCAM might be used as a surface marker to distinguish two groups of myoblasts prior to cell fusion.

To confirm the functional differences between two groups of myoblasts, we used both PSA-NCAM and NCAM antibodies for MACS-based cell sorting (Fig. 5). Our results showed that the PSA-NCAM positive myoblasts expressed higher levels of differentiation-related factors such as MYOG and exhibited faster myotube formation than the PSA-NCAM negative cells. This finding indicates that PSA-NCAM can serve as a surface marker to distinguish between proliferating and differentiated myoblasts.

To date, many signaling factors have been proposed to be involved in myoblast differentiation; however, most studies have evaluated the extent of differentiation based on myotube formation assay, making it difficult to separate the two stages of differentiation: phenotypic change and cell fusion. In contrast, the method established in this study allows for the detection of phenotypic changes prior to cell fusion, providing a higher-resolution methodology for elucidating the mechanisms of skeletal muscle differentiation.

To comprehensively capture this phenotypic change, we also conducted an exploratory RNA-sequencing analysis. PSA-NCAM positive and negative fractions were sorted by fluorescence-activated cell sorting (FACS) and compared them to publicly available single-cell RNA-sequencing data (Fig. S2-S5). First, using a similarity score based on PCA, we identified corresponding cell populations in clinical skeletal muscle samples, finding that both fractions showed high similarity to Muscle Satellite Cells (MuSCs) (Fig. S2) (De Micheli et al., 2020; Fitzgerald et al., 2023). On the other hand, when we concatenated clinical skeletal muscle data with cultured myoblast data (Wang et al., 2022), we found that both fractions were similar to cultured myoblasts group, though with subtle differences in distribution (Fig. S3-S4). Based on this distribution difference, trajectory analysis identified 12 genes that were associated with the transition from PSA-NCAM negative to positive states (Fig. S5). Gene enrichment analysis (Han et al., 2018; Zhou et al., 2019) showed enrichment in cell cycle-related terms and transcription factors, though the relationship with NCAM polysialylation remains a topic for future investigation.

In conclusion, we found that the state of NCAM in human skeletal muscle cells changes with differentiation and that this change can be identified using PSA-NCAM antibody. This discovery contributes to a more detailed understanding of skeletal muscle differentiation and regeneration mechanisms.

## Materials and methods

### Preparation of human skeletal muscle cells

Human (Homo sapiens Linnaeus, 1758) skeletal muscle cells (hSkMCs) were prepared from commercially available skeletal myoblasts (Lonza, Basel, Switzerland). The cells were expanded through two or three passages in the growth medium prepared by adding 10 ng/mL fibroblast growth factor-2 (FGF-2; Thermo Fisher Scientific, Waltham, MA, USA) and 125 ng/mL iMatrix-221 (Matrixome Inc., Osaka, Japan) to SkGM-2 medium (Lonza). The expanded cells were cryo-preserved in aliquots. Each experiment used cells from at least three different donor lots.

### Serum-free differentiation culture in avoiding fusion

Human skeletal muscle cells were plated in 100 mm tissue culture dishes at 2,500 cells/cm^2^ and cultured three days in serum-free differentiation medium prepared by supplementing 1 mg/mL recombinant human serum albumin (Nacalai Tesque, Kyoto, Japan), 1/100 ITS-G solution (Thermo Fisher Scientific), 500 ng/mL iMatrix-221, 100 U/mL Penicillin and 100 U/mL Streptomycin to high-glucose Dulbecco’s Modified Eagle Medium (FUJIFILM Wako Pure Chemical, Osaka, Japan) with the following growth factors and inhibitors added depending on the purpose of each experiment: 10 ng/mL FGF-2, 10 ng/mL epidermal growth factor (EGF; Miltenyi Biotec, Tokyo, Japan), 1 ng/mL transforming growth factor β1 (TGF-β1; Miltenyi Biotec), 1 μM SB505124 (Abcam, Cambridge, UK), 10 μM U0126 (FUJIFILM Wako Pure Chemical), 2 μM PD184352 (Cayman Chemical, Ann Arbor, MI, USA), 50 μM LY-294002 (FUJIFILM Wako Pure Chemical), 10 μM SB203580 (Adipogen Life Sciences, San Diego, CA, USA), 0.3 μM BIRB796 (Tocris Bioscience, Bristol, UK), and 10 μM SP600125 (Cayman Chemical).

### Flow cytometry

Multicolor flow cytometric analysis was performed for human skeletal muscle cell populations. When the primary antibodies were applied before fixation, the cells were harvested by Trypsin-EDTA (FUJIFILM Wako Pure Chemical) and treated by 1/400 volume primary antibody in serum-free medium for 30 min. After washing the primary antibodies, the cells were treated by 1/400 volume secondary antibodies for 30 min and washed. Then, the cells were fixed by 1% paraformaldehyde aqueous solution overnight in 4°C. When all primary antibodies were applied after fixation, the cells were fixed immediately after harvest. After fixation, washed cells were treated with (additional) primary antibody for 30 min with 0.1% Triton X-100 (Merck, Darmstadt, Germany) if necessary. After washing, the cells were treated with secondary antibodies and 0.5 μg/mL 4’,6-diamidino-2-phenylindole (DAPI; Thermo Fisher Scientific) for 30 min. Then, the cells were subjected to flow cytometric analysis using a Gallios flow cytometer (Beckman Coulter, Brea, CA, USA). All staining procedures except for fixation were conducted at room temperature. Phosphate buffered saline (PBS) with 1% Blocking One Solution (Nacalai Tesque) was used for washing and as the buffer to dilute antibodies.

Primary antibodies we used were: mouse monoclonal anti-NCAM(CD56) antibody (clone: MEM-188; BioLegend, San Diego, CA, USA), mouse monoclonal anti-desmin antibody (clone: D9; Thermo Fisher Scientific), rabbit recombinant monoclonal anti-desmin antibody (clone: RM234; Thermo Fisher Scientific), and mouse monoclonal anti-polysialylated NCAM (PSA-NCAM) antibody (clone: 12E3; Thermo Fisher Scientific). Secondary antibodies were: goat anti-mouse IgG1 antibody conjugated with Alexa Fluor 647 (Thermo Fisher Scientific), goat anti-mouse IgG2a antibody conjugated with Alexa Fluor 488 (Thermo Fisher Scientific), goat anti-mouse IgG antibody conjugated with Alexa Fluor plus 488 (Thermo Fisher Scientific), goat anti-rabbit IgG antibody conjugated with Alexa Fluor 647 (Thermo Fisher Scientific).

### Magnetic-activated cell sorting (MACS)

To fraction the PSA-NCAM positive myoblasts, PSA-NCAM negative myoblasts, and other cells (including non-myogenic cells), we exploited MACS technique. First, PSA-NCAM antibody (mouse IgM) was applied to the harvested cell suspension at 1/1000 volume for 30 min in 4°C. After washing, the cells were treated with magnetic beads conjugated anti-mouse IgM antibody (Miltenyi Biotec) for 30 min in 4°C. Then, the cells were separated using a magnetic separation column (Large Cell Columns; Miltenyi Biotec). The positively-selected fraction was stored as PSA-NCAM(+) cells. The flowthrough was further treated with magnetic beads conjugated anti-NCAM antibody (Miltenyi Biotec) for 30 min in 4°C and separated by another column. The positively-selected fraction and the flowthrough were stored as PSA-NCAM(-) cells and the flowthrough cells, respectively. These three fractions were subjected to flow cytometry, quantitative PCR or tube formation assay.

### Tube formation assay

To assess the extent of differentiation of myoblasts and tube formation, we performed differentiation culture and immunofluorescence imaging. The sorted cells were plated in Matrigel (1/100; growth factor reduced; Corning, Corning, NY, USA)-coated cell culture plates at 5×10^4^ cells/cm^2^ with differentiation medium. The cells were fixed with paraformaldehyde aqueous solution and washed by PBS. Then, mouse monoclonal anti-MYH4 antibody (clone: MF20; Thermo Fisher Scientific) and rabbit recombinant monoclonal anti-desmin antibody were applied at 1/1000 volume and treated for 1 h. After washing, the cells were further treated for 1 h with 1/1000 goat anti-mouse IgG antibody conjugated with Alexa Fluor plus 488 (Thermo Fisher Scientific) and 1/1000 goat anti-rabbit IgG antibody conjugated with Alexa Fluor 647, 1/1000 Alexa Fluor 568 conjugated Phalloidin (Thermo Fisher Scientific), and 0.2 μg/mL DAPI. Following washing, the fluorescent images were acquired using a laser scanning confocal microscope (Olympus, Tokyo, Japan) or ImageXpress Imaging System (Molecular Devices, San Jose, CA, USA). All staining procedure except for fixation was conducted in room temperature. PBS with 1% Blocking One Solution and 0.1% Triton X-100 was used for the buffer to dilute antibodies.

### Analysis of immunofluorescence images

The nuclei of all cells, desmin positive cells and MYH4 positive cells were identified on immunofluorescence images (16 images per a condition) using ImageJ software (public domain). Then, desmin positive ratio (myoblast purity) and MYH4 positive ratio in desmin positive population (tube forming ratio) were calculated.

### Gene expression analysis

From the MACS sorted cells, total RNA was extracted using the RNeasy plus Mini Kit (Qiagen, Venlo, Netherlands) according to the manufacturer’s instructions. cDNA was synthesized from the RNA using a RT-RamDA cDNA Synthesis Kit (TOYOBO, Osaka, Japan). Subsequently, gene expression was analyzed through TaqMan RT-PCR assay on a ViiA 7 Real-Time PCR System (Thermo Fisher Scientific). TaqMan probes for the target genes can be found in Supplementary Table S1 online. For the calculation of ΔCt, the geometric mean (arithmetic mean of Ct values) of the expression of three genes, TBP, RPLP0, and RPS17 was used as an endogenous control. These genes were determined by the geNorm method (Jo Vandesompele, Katleen De Preter, Filip Pattyn, Bruce Poppe,Nadine Van Roy, Anne De Paepe, 2002) from housekeeping genes in the TaqMan array human endogenous control plate (Thermo Fisher Scientific).

### Statistical analysis

Statistical analysis was conducted using two-way ANOVA and paired Student’s t-test. The results are presented on each figure.

## Acknowledgements

The authors thank the TWIns core facilities for their important contributions to this work.

## Data and resource availability

All relevant data and resource can be found within the article and its supplementary information.

## Author Contributions

T.K. designed and performed experiments, analyzed data. T.S. provided advice on research design and data interpretation. T.K. and T.S wrote the manuscript.

## Supplementary Data

**Table S1.** TaqMan probes for qPCR analysis.

| Target gene | Assay ID |
| --- | --- |
| ACTA1 | Hs00559403_m1 |
| TNNI2 | Hs00268536_g1 |
| TNNT1 | Hs00162848_m1 |
| TNNT3 | Hs00952980_m1 |
| MYH2 | Hs00430042_m1 |
| MYH3 | Hs01074230_m1 |
| MYH7 | Hs01110632_m1 |
| MYMK | Hs01368616_m1 |
| NCAM1 | Hs00941830_m1 |
| MEF2C | Hs00231149_m1 |
| MYOD1 | Hs00159528_m1 |
| MYOG | Hs01072232_m1 |
| TBP | Hs99999910_m1 |
| RPLP0 | Hs99999902_m1 |
| RPS17L | Hs00734303_g1 |

**Fig. S1.**
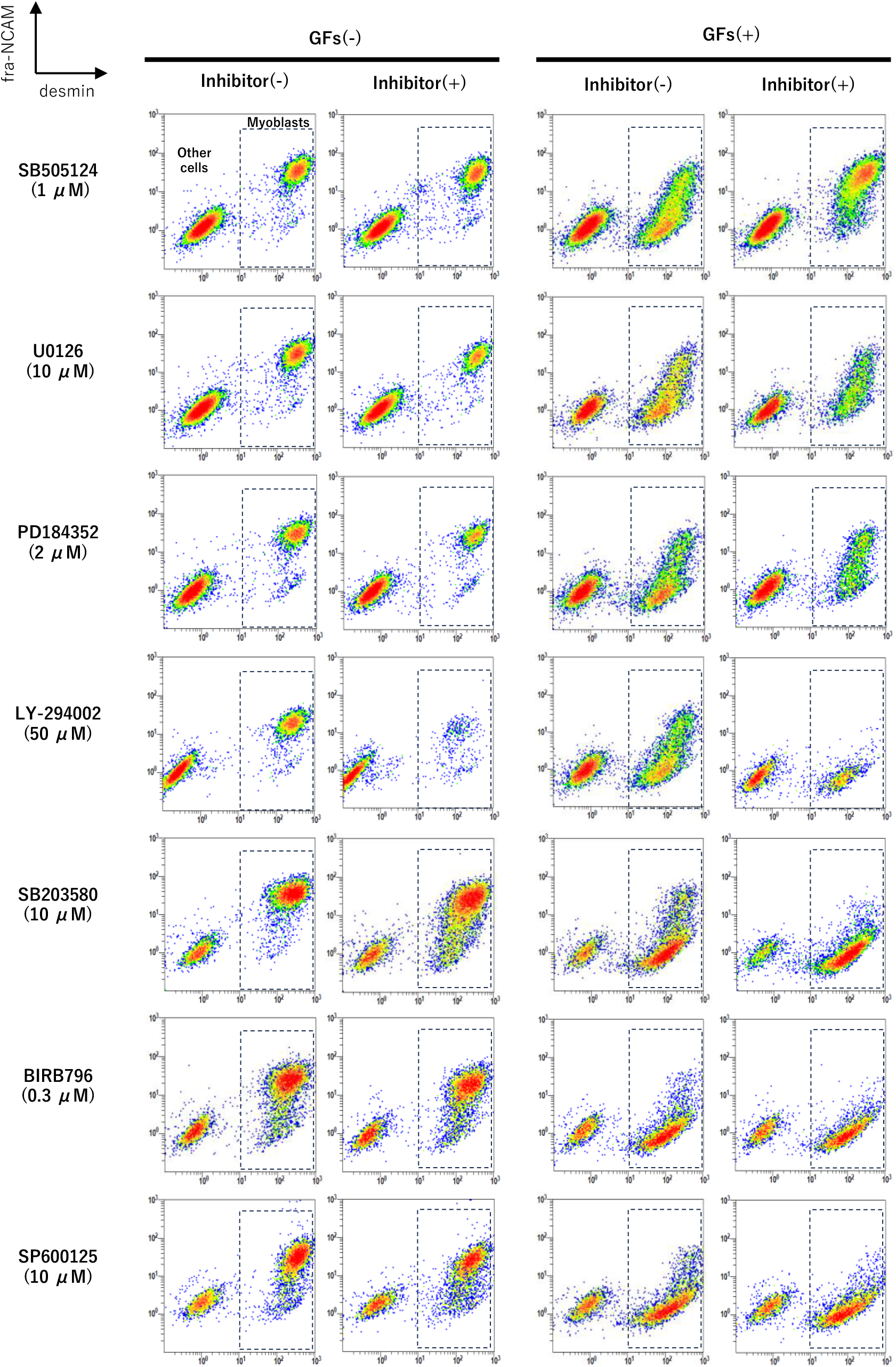
Representative scatter grams for signal inhibitors. hSkMCs were cultured in serum-free medium for 3 days with or without signal inhibitors under the conditions with or without growth factors (GFs: 10 ng/mL FGF-2, 10 ng/mL EGF and 1 ng/mL TGF-β1).

**Fig. S2.**
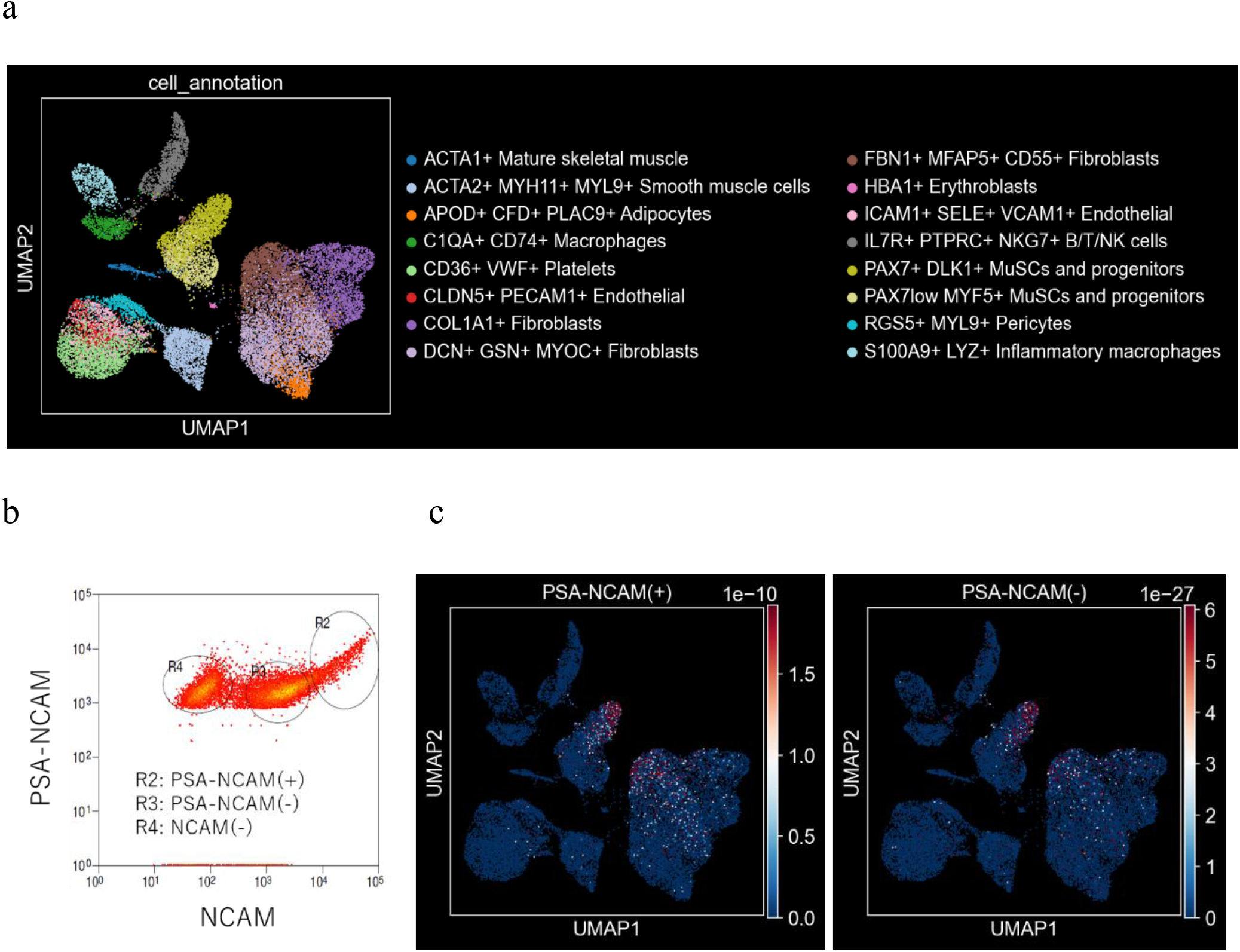
Similarity estimation on single-cell RNA-sequencing data of human skeletal muscle tissue biopsies. a) Cell annotations for cells in skeletal muscle tissue biopsy. Publicly available annotated single-cell RNA-sequencing (scRNA-seq) data (GSE143704) were re-analyzed. Downloaded raw count data were analyzed using R, with regression and normalization performed with SCTransform on feature counts, proportions of mitochondrial-related gene expression, and cell cycle scores. MuSCs: Muscle Satellite Cells. b) Fluorescence-activated cell sorting (FACS) of cultured human skeletal muscle myoblasts by using antibodies for PSA-NCAM and NCAM. c) Similarity scores for the conventional RNA-sequencing (RNA-seq) data of the PSA-NCAM positive and negative cells obtained by FACS. For RNA-seq and scRNA-seq data, all overlapping genes were extracted, normalized, and scaled to a standard normal distribution for each gene without regressing out. For scRNA-seq data, 2000 highly variable genes were extracted, and PCA was calculated. PCA was also calculated for the RNA-seq data using the same basis vectors, and the similarity was calculated by applying a Gaussian kernel to the Euclidean distance in PCA space.

**Fig. S3.**
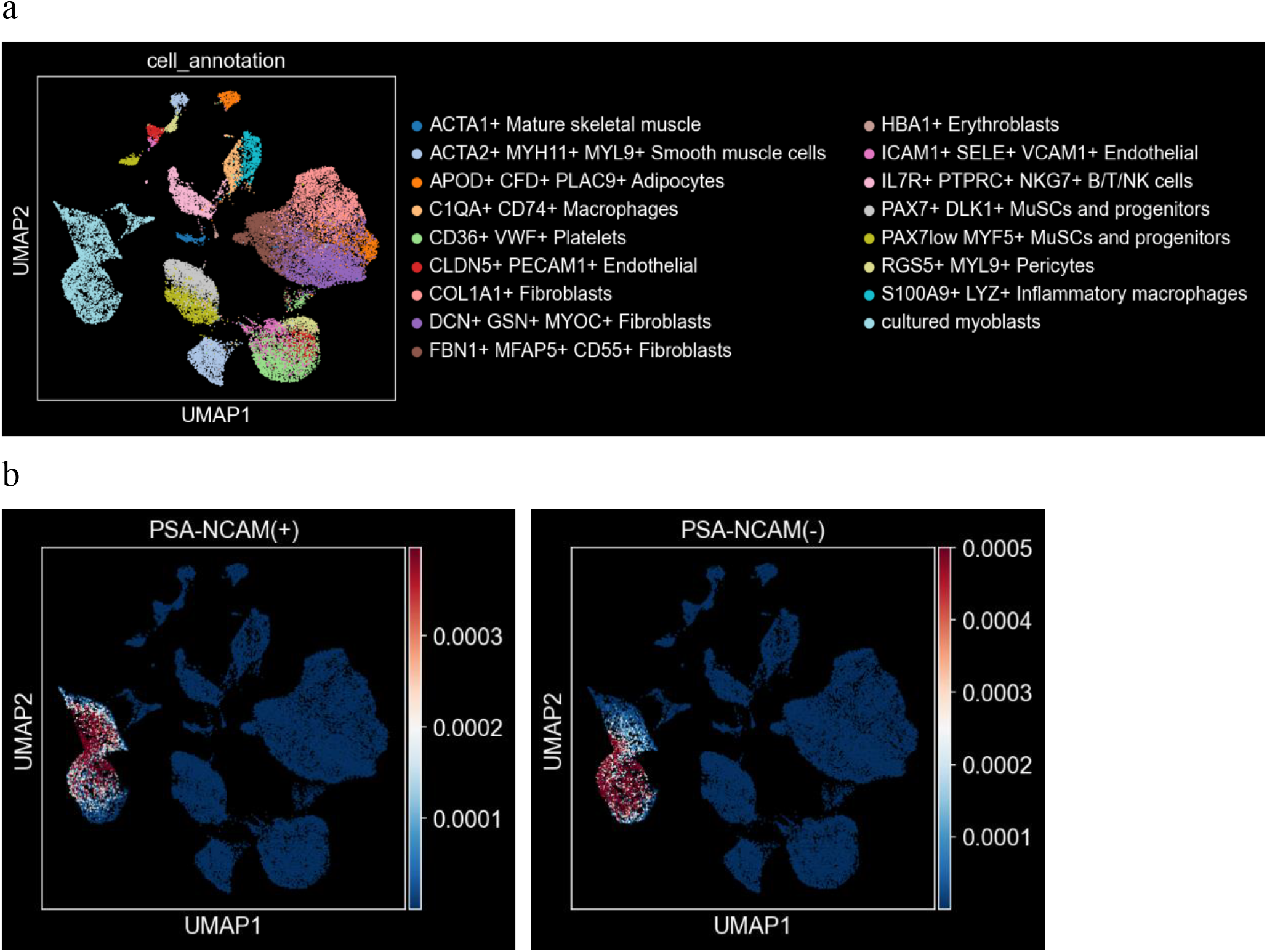
Similarity estimation on the combined single-cell RNA-sequencing data of both human skeletal muscle tissue biopsies and cultured myoblasts. a) Cell annotations. Single-cell RNA-sequencing (scRNA-seq) data of human skeletal muscle biopsies (GSE143704) and in vitro cultured myoblasts (GSE188215, group ‘2D’) were combined using SCTransform. MuSCs: Muscle Satellite Cells. b) Similarity scores for the conventional RNA-sequencing (RNA-seq) data of the PSA-NCAM positive and negative cells obtained by FACS. For RNA-seq and scRNA-seq data, all overlapped genes were extracted, normalized, and scaled to a standard normal distribution for each gene without regressing out. For scRNA-seq data, 2000 highly variable genes were extracted, and PCA was calculated. PCA was also calculated for the RNA-seq data using the same basis vectors, and the similarity was calculated by applying a Gaussian kernel to the Euclidean distance in PCA space.

**Fig. S4.**
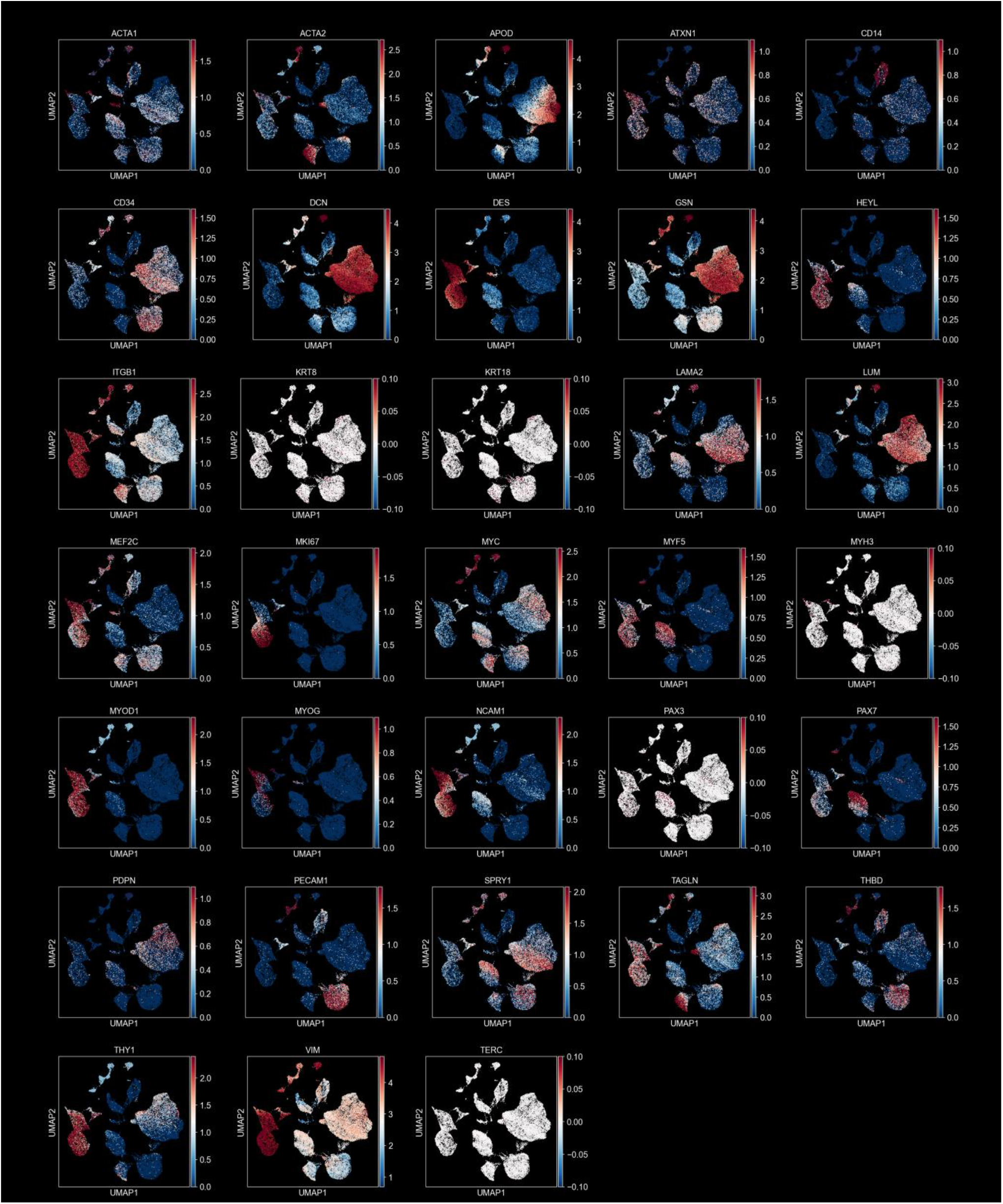
Gene Expression on the combined RNA-sequencing data of human skeletal muscle tissue biopsies and cultured myoblasts. a) Gene expression on the combined single-cell RNA-sequencing (scRNA-seq) data of human skeletal muscle biopsies (GSE143704) and in-vitro cultured myoblasts (GSE188215, group ‘2D’).

**Fig. S5.**
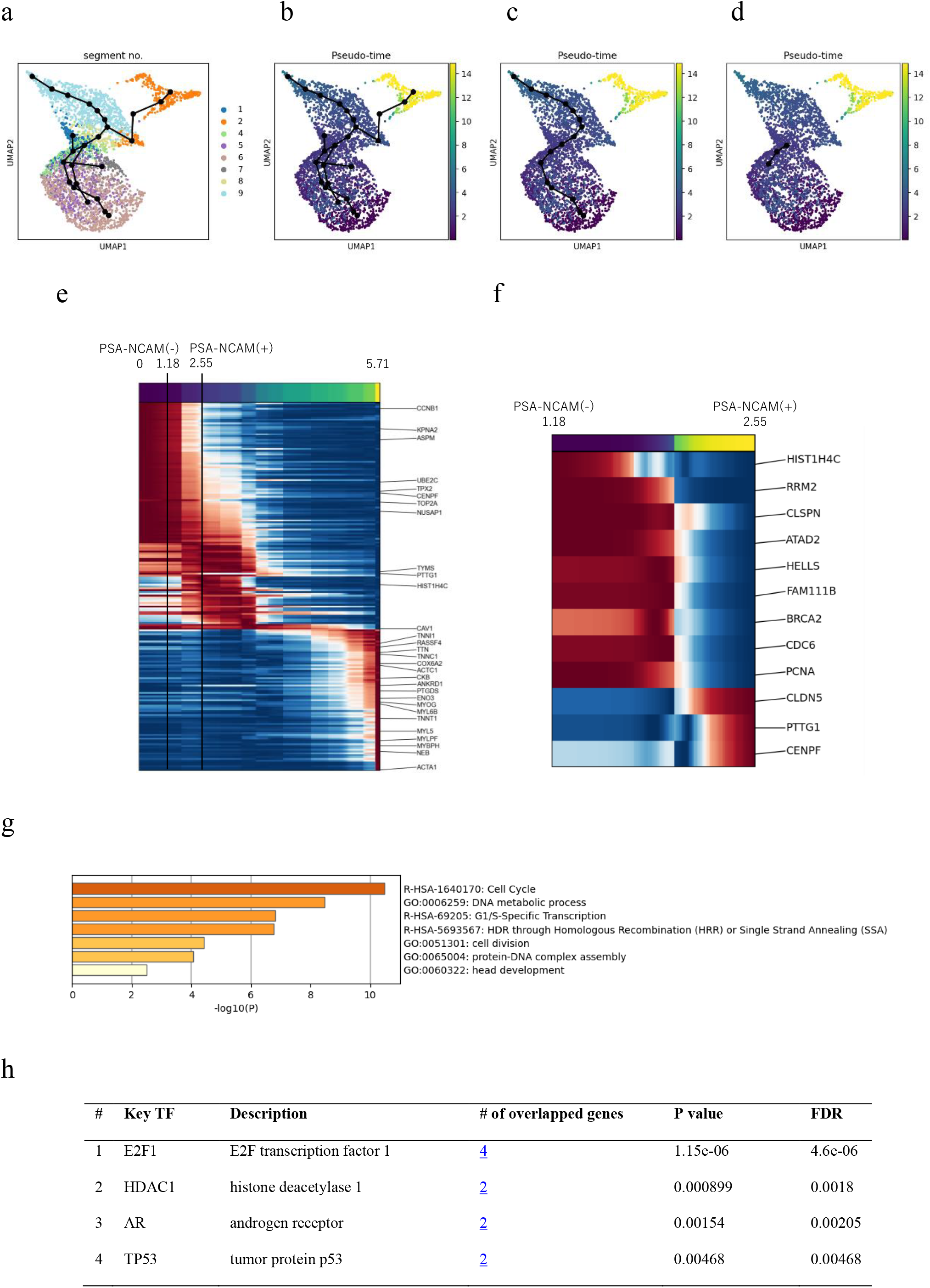
Trajectory analysis between PSA-NCAM(-) and PSA-NCAM(+) cells. a) The population of ‘cultured myoblast’ was extracted from the data of Fig.S3, and a principal tree was generated (python ‘scfates’ package) based on the multiscale diffusion map for PCA (python ‘palantir’ package). b) Pseudo-time analysis was performed using the tip point that appeared to be most proliferating (see MKI67 expression in Fig.S4) as the root. c) The probable, unbranched path from proliferative to differentiated myoblasts was chosen. d) The part of the path from PSA-NCAM(-) to PSA-NCAM(+) cells defined by each estimated pseudo-time. e) The set of genes significantly associated with the trajectory in panel c. X-axis represents pseudo-time. f) The set of genes significantly associated with the trajectory in panel d. X-axis represents pseudo-time. g) ‘Pathway and Process Enrichment Analysis’ in Metascape (https://metascape.org/) for the gene list of panel f. h) Candidate transcription factor analysis for the genes in panel f using TRRUST version 2 (https://www.grnpedia.org/trrust/).

